# Reward Circuit Local Field Potential Modulations Precede Risk Taking

**DOI:** 10.1101/2024.04.10.588629

**Authors:** Natasha C. Hughes, Helen Qian, Michael Zargari, Zixiang Zhao, Balbir Singh, Zhengyang Wang, Jenna N. Fulton, Graham W. Johnson, Rui Li, Benoit M. Dawant, Dario J Englot, Christos Constantinidis, Shawniqua Williams Roberson, Sarah K. Bick

**Author notes:** **Correspondence to:** Sarah Bick, MD, Department of Neurosurgery, Vanderbilt University Medical Center, 1500 21^st^ Ave South, Nashville, TN 37212.

## Abstract

Risk taking behavior is a symptom of multiple neuropsychiatric disorders and often lacks effective treatments. Reward circuitry regions including the amygdala, orbitofrontal cortex, insula, and anterior cingulate have been implicated in risk-taking by neuroimaging studies. Electrophysiological activity associated with risk taking in these regions is not well understood in humans. Further characterizing the neural signalling that underlies risk-taking may provide therapeutic insight into disorders associated with risk-taking.

Eleven patients with pharmacoresistant epilepsy who underwent stereotactic electroencephalography with electrodes in the amygdala, orbitofrontal cortex, insula, and/or anterior cingulate participated. Patients participated in a gambling task where they wagered on a visible playing card being higher than a hidden card, betting $5 or $20 on this outcome, while local field potentials were recorded from implanted electrodes. We used cluster-based permutation testing to identify reward prediction error signals by comparing oscillatory power following unexpected and expected rewards. We also used cluster-based permutation testing to compare power preceding high and low bets in high-risk (<50% chance of winning) trials and two-way ANOVA with bet and risk level to identify signals associated with risky, risk averse, and optimized decisions. We used linear mixed effects models to evaluate the relationship between reward prediction error and risky decision signals across trials, and a linear regression model for associations between risky decision signal power and Barratt Impulsiveness Scale scores for each patient.

Reward prediction error signals were identified in the amygdala (p=0.0066), anterior cingulate (p=0.0092), and orbitofrontal cortex (p=6.0E-4, p=4.0E-4). Risky decisions were predicted by increased oscillatory power in high-gamma frequency range during card presentation in the orbitofrontal cortex (p=0.0022), and by increased power following bet cue presentation across the theta-to-beta range in the orbitofrontal cortex (*p*=0.0022), high-gamma in the anterior cingulate (*p*=0.0004), and high-gamma in the insula (*p*=0.0014). Risk averse decisions were predicted by decreased orbitofrontal cortex gamma power (*p*=2.0E-4). Optimized decisions that maximized earnings were preceded by decreases within the theta to beta range in orbitofrontal cortex (*p*=2.0E-4), broad frequencies in amygdala (*p*=2.0E-4), and theta to low-gamma in insula (*p*=4.0E-4). Insula risky decision power was associated with orbitofrontal cortex high-gamma reward prediction error signal (*p*=0.0048) and with patient impulsivity (*p*=0.00478).

Our findings identify and help characterize reward circuitry activity predictive of risk-taking in humans. These findings may serve as potential biomarkers to inform the development of novel treatment strategies such as closed loop neuromodulation for disorders of risk taking.

## Introduction

Evaluating risks and benefits of available options is critical to our ability to make day-to-day decisions. Increased propensity for risk-taking can be present in many psychiatric and neurological diseases including substance use disorder,^1–3^ bipolar disorder,^4^ frontotemporal dementia,^5^ and Parkinson’s disease.^6–8^ Deficits in the ability to evaluate and avoid risk can contribute to significant morbidity for patients with these diseases, leading to unfavourable choices impacting their quality of life and increasing overall risk of death.^9–12^ Unfortunately, this pathologic tendency towards risk-taking is not well-managed by current treatment regimes.^13,14^ Understanding neural activity changes associated with risk-taking decisions is critical for better understanding the underlying mechanisms that give rise to these pathologies and developing targeted treatments.

Interactions between reward circuitry regions such as the orbitofrontal cortex (OFC), amygdala, insula, and anterior cingulate cortex (ACC) have been implicated in processing and evaluating rewards and thus impacting decision-making.^15–17^ Reward processing is performed primarily by the mesolimbic system, with the ventral tegmental area thought to play a major role in directing reward-related responses and communicating reward-related information directly and indirectly to other limbic regions such as the prefrontal cortex, amygdala, cingulate, and insula.^18–20^ Moreover, reward processing signals have been shown to be linked to risk-taking.^21,22^ Reward prediction error signals (RPE) are signals associated with the difference between expected and received reward coded by dopaminergic neurons, and these signals have been suggested to modulate future risk-taking.^23^ In rats, an artificial positive RPE signal created by ventral tegmental area stimulation following nonrewarded choices has been shown to increase future risky choices.^24^ Imaging studies have also suggested that activity in these regions may predict risk taking decisions. 18-fluorodeoxyglucose positron emission tomography in patients with Parkinson’s disease showed that increased insula, OFC, and ACC metabolism is associated with increased impulsivity scores.^25^ Furthermore, human neuroimaging studies suggest that the insula,^26,27^ ACC,^28,29^ OFC,^28,30^ and amygdala^31–33^ modulate their activity prior to and during risky decision making and while evaluating risk.

A limitation of current understanding of reward circuit signalling associated with risk taking behavior has been intrinsic limitations of the techniques used to measure neural activity changes. Neuroimaging studies are limited in temporal resolution due to hemodynamic response time and in spatial resolution due to signal-to-noise ratios and size of regions of interest.^34^ In contrast, electrophysiology techniques have exquisite temporal resolution and more directly measure neural activity, and thus overcome these limitations of functional imaging techniques. Scalp electroencephalography (EEG) has demonstrated that neural activity in prefrontal cortex regions predicts risk-taking behaviours in human subjects.^35–37^ However, scalp EEG approaches are limited in spatial resolution and in assessing deep brain regions, such as the ACC, insula, OFC, and amygdala. Intracranial EEG performed in human neurosurgical patients can overcome these limitations by directly recording from deep brain structures with excellent temporal and spatial resolution.

We used intracranial EEG to characterize neural oscillatory activity predictive of a risky decision. We recorded from the amygdala, ACC, insula, and OFC in patients with medically refractory epilepsy who as part of their regular clinical course received implanted depth electrodes in these regions. Subjects participated in a card-based gambling task and made decisions to take more or less risk. We then evaluated neural signals that occurred prior to these decisions. Finally, we evaluated reward related signals in these structures and their relationship to risk-taking signals. Evaluating how reward circuitry structures represent reward and predict subsequent risk-taking could provide insight into future therapeutic targets for disorders of risk-taking.

## Materials and methods

### Participants

We recruited eleven patients with medication resistant epilepsy who, as a part of their clinical evaluation, were implanted with stereotactic-electroencephalography (SEEG) electrodes to localize their seizure focus. Electrode implant locations for each patient were clinically decided based on preoperative hypotheses of site of seizure onset. Following implantation, patients were admitted to the epilepsy monitoring unit where they were monitored for seizures and their anti-seizure medications were gradually weaned. All subjects provided written informed consent prior to participating in this study and this study was approved by the Institutional Review Board of Vanderbilt University Medical Center (Nashville, TN, IRB #211037).

The day prior to participating in the behavioural task, patients completed the Barratt Impulsiveness Scale (BIS-11). The BIS-11 consists of 30 items graded on a four-point scale (1-4) to assess for general impulsivity. Raw scores range from 30-120 with greater scores indicating greater impulsivity; scores between 52-71 are thought to be as within normal limits, with >71 indicating highly impulsive and <52 indicating over-controlled.^38^ One patient did not complete the BIS-11.

### Electrode Localization

Each participant underwent placement of multiple sEEG leads each containing 8-16 contacts spaced 2.5-4.3 mm center-to-center (PMT corporation, Chanhassen, MN). The contact coordinates were identified by superimposing preoperative MRI and postoperative CT images using CRAnial Vault Explorer (CRAVE) software.^39^ Location of these coordinates within specific brain regions was then identified using the Desikan-Killiany Atlas^40^ using FreeSurfer’s cortical parcellation and subcortical segmentation procedure.^41,42^ An epileptologist (co-author SWR) manually verified these locations based on visual inspection of the co-registered MRI postsurgical CT, and baseline EEG timeseries.

Electrodes determined to be within the OFC, amygdala, insula, and ACC were included for analysis. Channels determined by the clinical team and by a clinical epileptologist (co-author SWR) to be within the patient’s seizure onset zone were excluded from analysis. Of the eleven participants, ten had OFC electrodes, six had ACC electrodes, ten had amygdala electrodes, and ten had insula electrodes that were included in the final analysis.

### Experimental Design

Participants performed a gambling task using a portable tablet and keyboard while continuous local field potentials were recorded from sEEG electrodes using the clinical recording system (Natus). The task was presented on a portable 13-inch tablet (Microsoft Surface, Redmond WA) using Psychophysics Toolbox Version 3 core and MATLAB 2022 (MathWorks, Natick, MA).^43^ This experiment was performed between 1-2 days post-operatively while patients were in the epilepsy monitoring unit of the hospital in specially-equipped rooms with continuous EEG and video recordings. Digital pulses were sent to the trigger channel of the EEG data acquisition system in conjunction with task events in order to synchronize task events with EEG recordings.^44^

Patients were instructed to maximize their winnings in a gambling task. In this task, they were shown a card from a deck made up of the cards 2, 4, 6, 8, and 10 and were instructed to place a wager as to whether their card was higher than a hidden card. They could bet either high ($20) or low ($5). Patients could either win, lose, or tie depending on the hidden card. Each trial, there were 5 behavioural events that occurred in sequence: 1) patient card presentation, 2) bet cue presentation, 3) patient response, 4) opponent card presentation, and 5) result reveal (**Figure 1A**). The placement of $5 and $20 on right or left sides of the screen during bet cue presentation indicating which button the subject should press to record the respective bet was randomized each trial to control for directional effects. Before participation in study recordings, a member of the experimental team explained the task to the subject and the patient participated in a practice block for at least 10 trials, or until they felt comfortable with the task. These practice blocks were not recorded. After this, patients participated in 2 blocks of the gambling task of 75 trials each. Trials in which the patient took longer than 10 seconds to respond to the bet cue were excluded from analysis.

**Figure 1.**
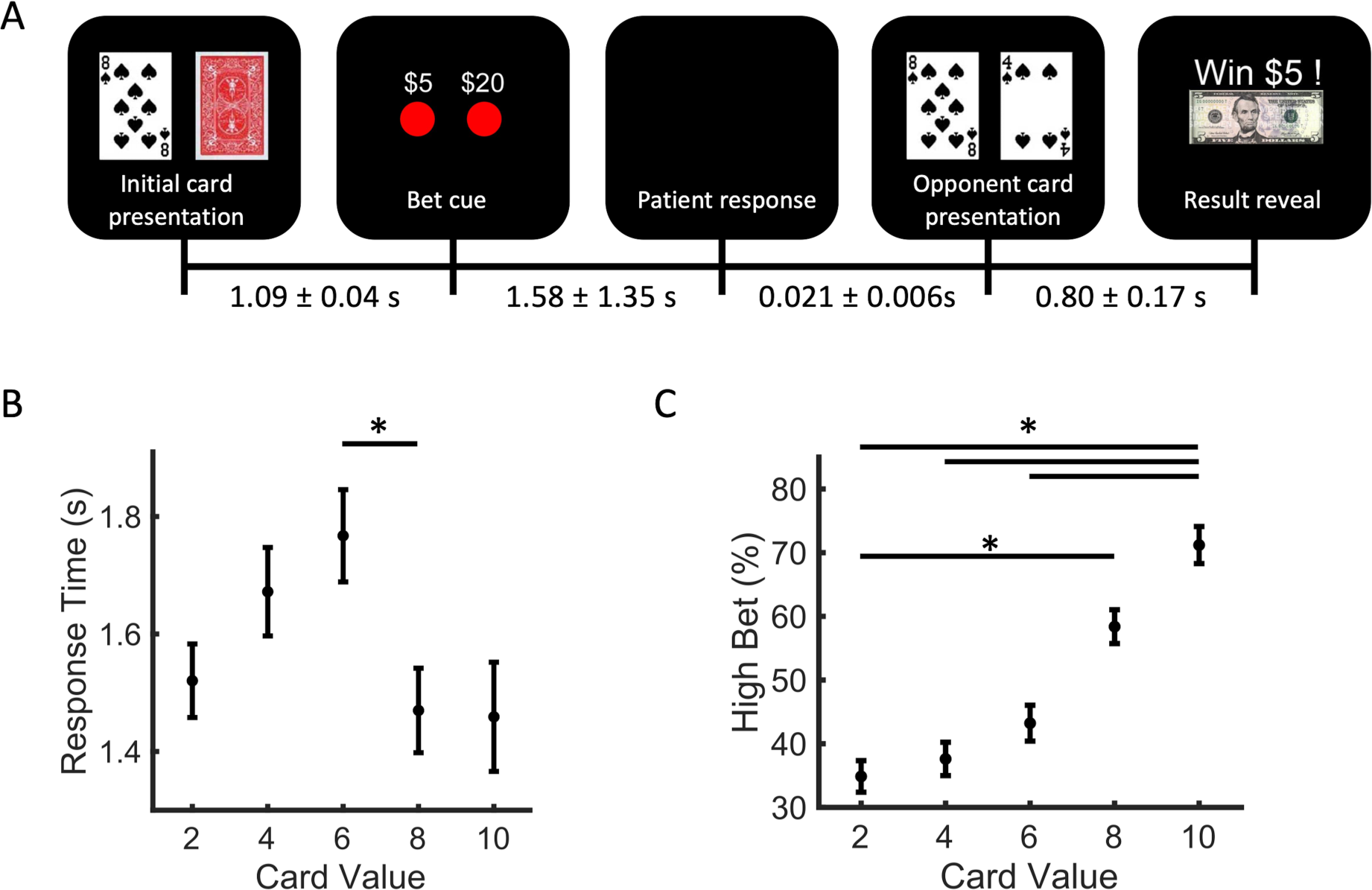
Gambling task and behavioural data. (A) Behavioural task events and timing between events in mean seconds ± standard deviation. (B) Mean response time (time from bet cue presentation to patient response) for each patient card value. Error bars represent standard error. Response time for 6 cards was significantly longer than that for 8 cards. (C) Mean percent of trials on which subjects bet high ($20) for each patient card value. * indicates p < .05. Error bars represent standard error.

Trials were separated into “high risk” and “low risk” based on percent chance of winning. If a patient had less than or equal to 50% chance of winning, this was classified as high risk (a presented card of 6 or less) whereas if the patient had greater than 50% chance of winning, this was considered low risk (a presented card of 8 or greater).

### Electrophysiology Recording and Pre-Processing

SEEG recordings were collected using the Natus clinical data acquisition system (Natus, Middleton, WI) at a sampling rate of 512 Hz and were referenced to a scalp electrode contact. Digital signals sent from the task presentation tablet allowed alignment of SEEG recordings with task events including patient card presentation, bet cue presentation, patient response, opponent card presentation, and result reveal.

SEEG pre-processing was performed using MATLAB and FieldTrip MATLAB toolbox.^45^ SEEG data was filtered with a bandpass filter from 1-200Hz with padding and with a 60 Hz, 120 Hz, 180 Hz notch filter to remove line noise. Data was then re-referenced to the common average across all channels. All traces were manually inspected for channels or trials with significant artifacts which were excluded from further analysis.

For each trial and channel, time-frequency representations of power for each 2 Hz frequency interval within the range of 2-100Hz was computed using a Hanning window of 0.5 seconds that slides in steps of 0.01 seconds. Custom MATLAB code (MATLAB 2020b) was used for further analysis. Oscillatory power was Z-scored for each channel and frequency across each block to allow comparison across subjects and channels.

Processed SEEG data were segmented into epochs by alignment to the following behavioural events: (1) card presentation epochs, lasting from time of initial patient card presentation to 1000ms after, (2) bet cue epochs, lasting from time of presentation of the cue to bet until 2000ms afterwards, (3) result-locked epochs, lasting from time of result reveal until 1000ms after.

### Statistical Analysis

For behavioural analysis, a repeated measures ANOVA was used to identify relationships between card value and patient response times. Tukey’s post-hoc honest significant difference test was then used to identify significant differences between response times for each card value. A repeated measures ANOVA was used to determine if response times differed in trials where patients bet high compared to trials where patients bet low, controlling for each patient as a random variable. Similarly, a repeated measures ANOVA accounting for individual patient variability followed by Tukey’s post-hoc honest significant difference test was also used to determine relationship between average percent high bet across patients in card values and to compare responses for each presented card value.

For SEEG analysis, we used cluster-based permutation testing to identify oscillatory power changes associated with risk-taking behavior and reward signalling. First, we identified RPE signals using two-sided dependent cluster-based permutation testing (an alpha threshold of 0.05 and 5000 permutations) to compare mean power for trials in which expected reward (reward in a trial with >50% chance of winning) compared to unexpected reward (a reward in a trial with <50% chance of winning) was received during the result reveal epoch.^46^ Significant time frequency clusters were defined as those with *p* < 0.01 to correct for multiple structures examined. Secondly, we identified risk-related signals prior to a bet decision being made using this same procedure to compare mean power for high-bet high-risk to low-bet high-risk trials for both the card presentation and bet cue aligned epochs. Adjacent time frequency clusters with border points less than 0.05 seconds and 10 Hz apart and with more than 50% of the smaller cluster’s frequency values represented within the larger cluster were combined into a single cluster for further analysis.

For each identified time frequency cluster, we averaged z-scored power within the identified time and frequency ranges for trial types of interest on a per channel basis. To identify oscillatory power signals associated with risk, we examined mean z-scored power within identified clusters for each combination of bet and risk (high-bet high-risk, low-bet high-risk, high-bet low-risk, and low-bet low-risk) as well as each combination of bet and card value (high bet with a card value of 2, low bet with a card value of 2, etc.). These average z-scored power values for each condition were used in two-way ANOVAs to identify associations between cluster power, bet, and risk, and between cluster power, bet, and card value. Post-hoc Tukey’s honest significant difference was then used to determine significant differences in mean power between different conditions. We similarly calculated mean z-scored power within RPE-identified clusters for each combination of reward expectation and reward magnitude (unexpected rewarded $5, unexpected rewarded $20, expected rewarded $5, expected rewarded $20) and used two-way ANOVA to identify associations between reward expectation, reward magnitude, and cluster power. The Holm-Bonferroni method was used to correct two-way ANOVA and post-hoc Tukey’s statistics for multiple comparisons across the nine identified risk-associated signals and four RPE signals. This correction’s use is reported as an “adjusted” p-value.

To examine signals associated with risk, we used results from two-way ANOVA and Tukey’s honest significant difference to categorize identified clusters. We defined risk signals as clusters with significant effect of risk on two-way ANOVA, but without significant difference in mean power between high and low bets in the high-risk condition by Tukey’s post-hoc test. We also identified three risk and bet interaction signal types: risky decision signals, risk aversion signals, and optimal decision signals. Risky decision signals were clusters in which power predicted high-bets in the high-risk condition; these, in addition to being significantly different between high and low bets in the high-risk condition, were clusters whose average power was significantly modulated by either trial risk or bet-risk interaction in the two-way ANOVA. Risk aversion signals were clusters associated with bet and risk that had significant difference in power in the high-risk condition prior to the patient betting low relative to all other conditions (in other words, a signal associated with avoidance of the riskiest option). Lastly, optimal decision signals were clusters with mean power associated with bet and risk and with, by Tukey’s post-hoc test, greater average power prior to decisions that are “optimized” for winning the most amount of money. In other words, optimized decision signals had greater power in high-risk scenarios prior to betting low than prior to betting high, and had greater power in low-risk scenarios prior to betting high than prior to betting low.

To evaluate whether risky decision and RPE signals were related, we calculated average z-scored power within each risky decision and RPE cluster’s time and frequency boundaries for each high-bet high-risk trial that resulted in a reward. In order to examine the impact of RPE in all four brain regions on risky decision signals we limited this analysis to the five patients with leads in all four brain regions of interest. We then used linear mixed effects models, controlling for patient identity, with power of all RPE signals as predictors to evaluate the associations between these signals and each of the four risky decision signal (alpha=0.0125, corrected for multiple comparisons for 4 risky decision signals).

To determine whether risky decision neurophysiology signals were associated with impulsive behavior we evaluated the relationship between power in risky decision clusters and BIS-11 scores. For each risky decision signal, we calculated the average difference in Z-scored power between high-bet high-risk trials and low-bet high-risk trials for each subject. We used a linear regression model to determine if this difference in z-scored power was associated with BIS-11 scores. One patient without a recorded BIS-11 score was excluded from this analysis.

## Results

### Behavioural Results

Patient demographics, seizure foci, and BIS-11 scores are shown in **Supplemental Table 1**. The average BIS-11 score of these patients fell within the normal range at 68.9 +/− 11.3 and the average percent of high-risk trials in which a patient bet high was 39.1% (+/− 16.6%).

Average response time was 1.58 +/− 1.35 seconds (**Figure 1A**). We also evaluated whether response times were associated with card identity and found that response time and card value were significantly associated (*F*(4,1607)=3.18, *p*=0.023). Cards associated with greater uncertainty had longer response time, with the longest mean response time in trials with the card value of 6 (**Figure 1B**). Post-hoc testing demonstrated a significant difference between mean response times for card values of 6 and 8 *(p=*0.023), but differences between mean response times for other pairs of card values were not significant. Additionally, when adjusting for patient differences, patients were more likely to place a high bet when the card value, and thus chance of winning, was greater, indicating that patients used appropriate strategy to participate in the task (*F*(4,54)=7.07, *p*=0.0002) (**Figure 1C**). Though patient ID was associated with response times to the task (*F*(10,1601)=48.03, *p*=4.15E-7), there was no significant association between response times and high or low bets (*F*(1,1610)=0.94, *p*=0.35) and no interaction between patient ID and bet choice in impacting response time (*F*(10,1601)=1.11, *p*=0.35).

### Reward Prediction Error Signals

We examined whether our brain regions of interest had power changes associated with RPE. To do this we compared power during the result reveal epoch of the task between high-risk win (unexpected win) and low-risk win (expected win) trials. The OFC, ACC, and amygdala all had power changes associated with delivery of an unexpected reward, which we considered RPE signals (**Figure 2**). The OFC had two clusters that differed between unexpected and expected wins, one in high gamma 70-90 Hz frequency range from 0.01-0.47 seconds after result reveal that had increased power during unexpected rewards (*p*=6.0E-4), while the other was in theta-alpha 4-18 Hz frequency range from 0.34-0.97 seconds after result reveal with decreased power during unexpected rewards (*p*=4.0E-4) (**Figure 2A,E**). On two-way ANOVA average power within both clusters was associated with expectation of reward (*F*(1,526)=57.37, adjusted *p*=2.3E-11, and *F*(1,526)=41.15, adjusted *p*=7.1E-9 respectively); the higher frequency signal was not affected by reward magnitude (*F*(1,526)=0.01, adjusted *p*=1) whereas the lower frequency signal was associated with reward magnitude (*F*(1,526)=15.33, adjusted *p*=0.00043) and the interaction between reward magnitude and reward expectation (*F*(1,526)=11.64, adjusted *p*=0.0026). In ACC, there was a 56-64 Hz RPE signal that decreased in power 0.01-0.81 seconds following unexpected rewards (*p*=0.0092 for cluster; *F*(1,142)=12.96, adjusted *p*=0.00097 for reward expectation) (**Figure 2B,F**). The amygdala similarly had an RPE signal (*p*=0.0066) with decreased power within the 28-32 Hz frequency range 0.23-0.85 seconds following unexpected rewards (*p*=0.0066 for cluster; *F*(1,230)=19.17, adjusted *p*=0.0001 for reward expectation) (**Figure 2C,G**). Both ACC and amygdala signals were not associated with reward magnitude (*F*(1,142)=0.8, adjusted *p*=0.38 and *F*(1,230)=4.95, adjusted *p*=0.060 respectively). The insula did not have any significant RPE signals (**Figure 2D**).

**Figure 2.**
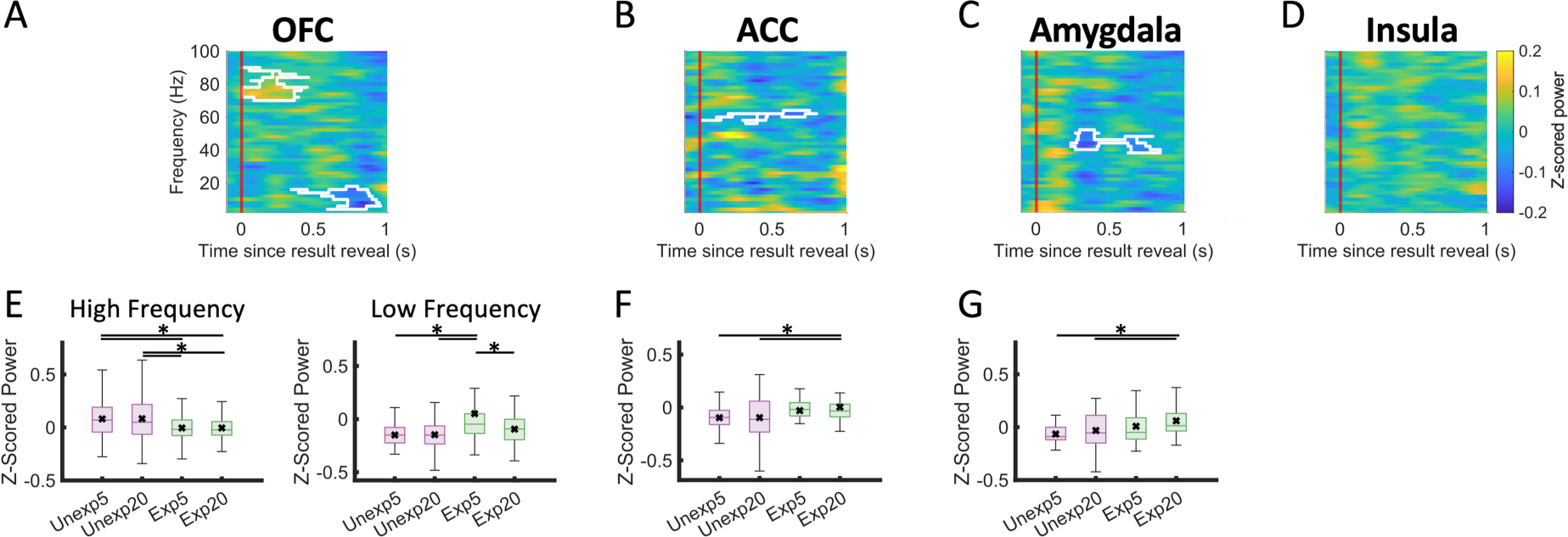
Reward prediction error (RPE) signals. Power spectra visualizations of difference in average power during result reveal between unexpected and expected wins. White outlines indicate time frequency clusters in which the difference between these two conditions is statistically significant. Red line (time=0s) indicates start of the result reveal epoch. This is visualized for the **(A)** orbitofrontal cortex (high frequency *p*=6.0E-4, low frequency *p*=4.0E-4), **(B)** anterior cingulate (p=.0092), **(C)** amygdala (p=.0066), and **(D)** insula. Below each power spectra is a box plot of the mean power within the significant cluster across the four conditions of reward expectation and reward magnitude for the orbitofrontal cortex **(E**), anterior cingulate **(F**), and the amygdala **(G)**. Significant difference between groups (adjusted p<0.05) is indicated with line and * above. Black x indicates mean, whereas horizontal line within box indicates median. OFC = orbitofrontal cortex, ACC = anterior cingulate cortex. Unexp5 = unexpected reward, rewarded $5, Unexp20 = unexpected reward, rewarded $20, Exp5 = expected reward, rewarded $5, Exp20 = expected reward, rewarded $20.

### Risk Signal

To identify oscillatory power changes associated with risk taking we used cluster-based permutation testing to identify time frequency clusters of power in high risk trials that were significantly different between conditions in which subjects ultimately bet high compared to those in which they bet low (**Figure 3**). We then used two-way ANOVAs to identify relationships between power within these clusters, risk level, and bet choice. One significant time frequency cluster within the OFC was associated with trial risk level but was not significantly different between high and low bets in the high-risk condition (adjusted *p*=1) (**Figures 3B, 4A,B**). This cluster had an increase in power within the 40-100Hz frequency range from 0.23 to 1.48 seconds after bet cue presentation during high risk trials (*p*<0.01). Power in this cluster was significantly associated with risk (*F*(1,526)=62.98, adjusted *p*=1.6e-12), bet (*F*(1,526)=10.7, adjusted *p*=0.0027), and the interaction between bet and risk (*F*(1,526)=6.60, adjusted *p*=0.023). This signal had significantly increased power in high-risk relative to the low-risk trials (adjusted *p*=<0.0001 suggesting this is a signal primarily associated with risk level (**Figure 4A,B**). Unlike in the high-risk condition, in the low-risk condition this signal did differentiate bet choice, with significantly lower power preceding low relative to high bets (adjusted *p*=5.8e-05) (**Figure 4B**).

**Figure 3.**
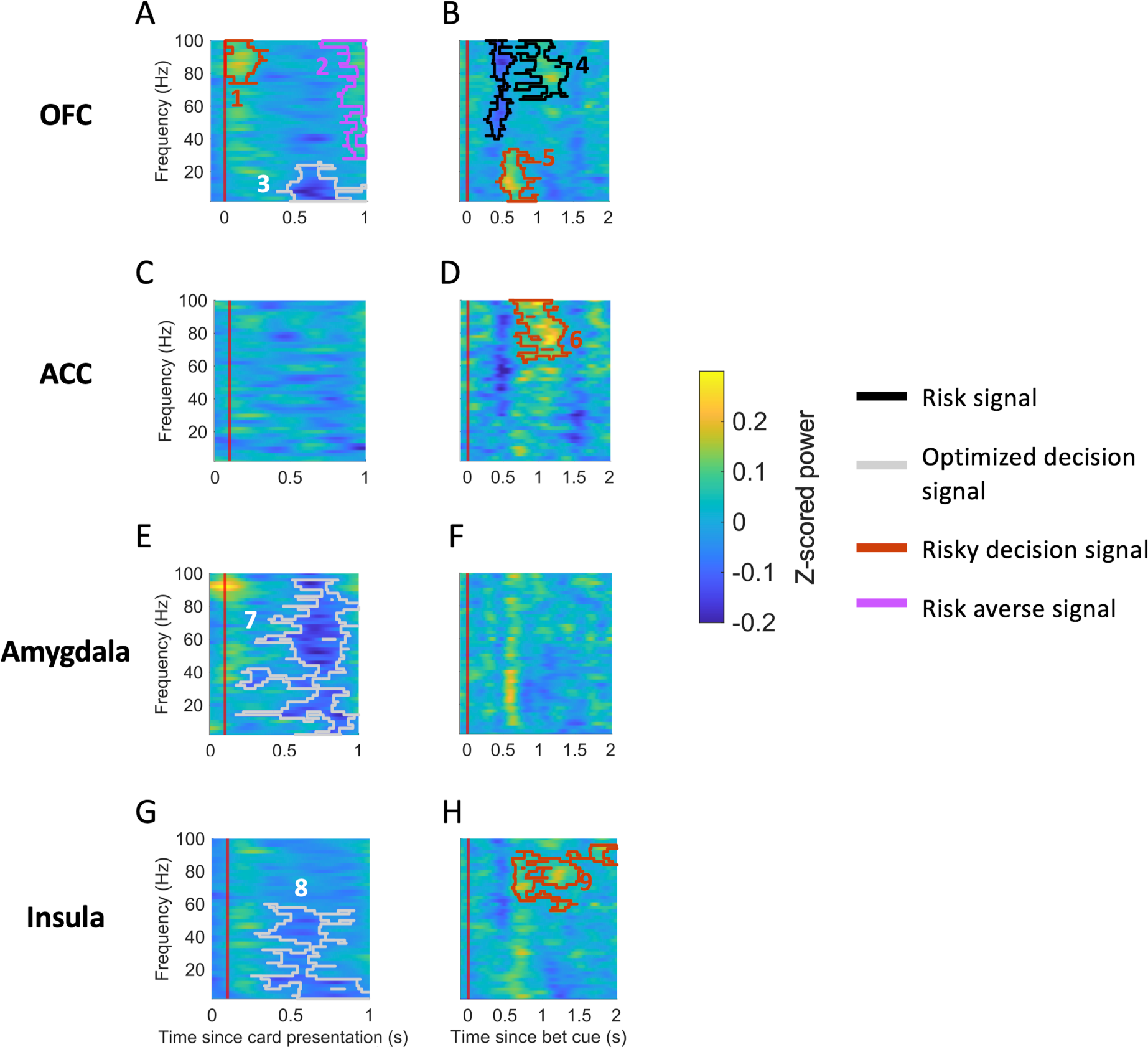
Risk related power signals. Power spectra visualizations of difference in average power during card presentation **(A,C,E,G)** and bet cue **(E,D,F,H)** between trials in which subjects subsequently bet high versus low in the high-risk condition. Outlined regions are significant clusters with colours corresponding to signal type as shown in legend. Red line (time = 0s) indicates start of alignment event. Orbitofrontal cortex signals are shown aligned by time since patient card presentation (Signals 1,2,3: p=.0022, p=2.0E-4, p=2.0E-4 respectively) **(A)** and aligned by bet cue (Signals 4,5: p<.01, p=.0022 respectively) **(B)**. Anterior cingulate cortex signals are shown aligned by card presentation **(C)** and by bet cue (Signal 6: p=4.0E-4) **(D)**. Amygdala signals are shown aligned by card presentation (Signal 7: p=2.0E-4) **(E)** and by bet cue **(F)**. Lastly, insula signals are shown aligned by card presentation (Signal 8: p=4.0E-4) **(G)** and by bet cue (Signal 9: p=.0014) **(H)**.

**Figure 4.**
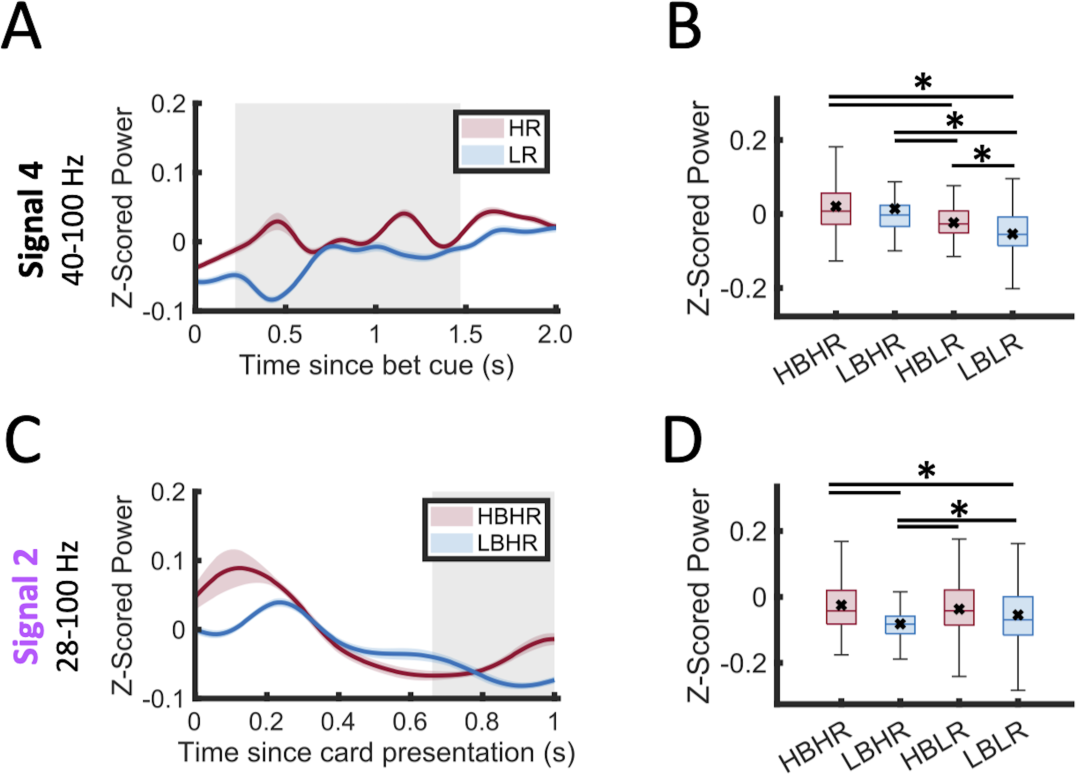
Orbitofrontal cortex risk and risk aversion signals. Average z-score power within cluster frequency range plotted over time since bet cue in **(A)** high risk, in red, and in low risk, in blue, conditions showing risk-related signal or **(C)** high bet high risk, in red, and low bet high risk, in blue, conditions showing risk aversion signal. Shaded bars indicate standard error. Grey box indicates time range of significant cluster. Box plot of the mean power within the significant cluster across the four conditions **(B, D)**. Significant difference between groups (adjusted p<0.05) is indicated with line and * above. Black x indicates mean, whereas horizontal line within box indicates median. (HB = high bet; LB = low bet; HR = high risk; LR = low risk).

### Signals Associated with Bet and Risk

All four studied regions had clusters whose power was associated with both bet and risk. In particular, we identified three types of bet and risk interaction signals: ones that predict aversion to a risky choice – a “risk aversion signal,” ones that predict risky choices – a “risky decision signal,” and finally ones that represent an optimized bet to maximize gains and minimize loss – an “optimized decision signal.”

### Risk Aversion Signals

We defined risk aversion signals as those that predicted low bets in high-risk trials, and found one OFC cluster associated with risk aversion. This was a 28-100Hz power decrease in low bet high risk trials compared to all other conditions that occurred 0.67-1.00 seconds after subject card presentation (*p*=2.0E-4 for cluster, *F*(1,526)=9.51 and adjusted *p*=0.01 for interaction of bet and risk) (**Figures 3A, 4C,D),** suggesting that a decrease in power within this range was associated with subsequent risk avoidance behavior.

### Risky Decision Signals

We defined risky decision signals as those that predicted high bets in high risk trials. The OFC, ACC, and insula had time frequency clusters whose power was significantly associated with subsequent risky decisions. The OFC had two clusters where increase in power was associated with a subsequent risky decision. The first was an increase in gamma range (74-100Hz) oscillatory power 0-0.31 seconds after patient card presentation (*p*=0.0022 for cluster, *F*(1,526)=7.48 and adjusted *p*=0.021 for interaction of bet and risk) (**Figure 3A**). A second, lower frequency (2-34 Hz) cluster was seen 0.4-1.04 seconds after bet cue presentation (*p*=0.0022 for cluster, *F*(1,526)=15.96 and adjusted *p*=5.4E-4 for bet and risk interaction) (**Figure 3B**). For both of these signals, mean power was significantly greater prior to high-bet decisions in high-risk conditions when compared to all other bet and risk level combinations (adjusted *p*<0.05 against all other risk-bet conditions) (**Figure 5 A-F**). The power within these clusters was also both significantly associated with card value and the bet-card interaction (adjusted *p*<0.001) (**Figure 5 C,F**). In general, for both of these clusters power prior to high bets was greater for lower value cards, suggesting modulation of this signal by risk level.

**Figure 5.**
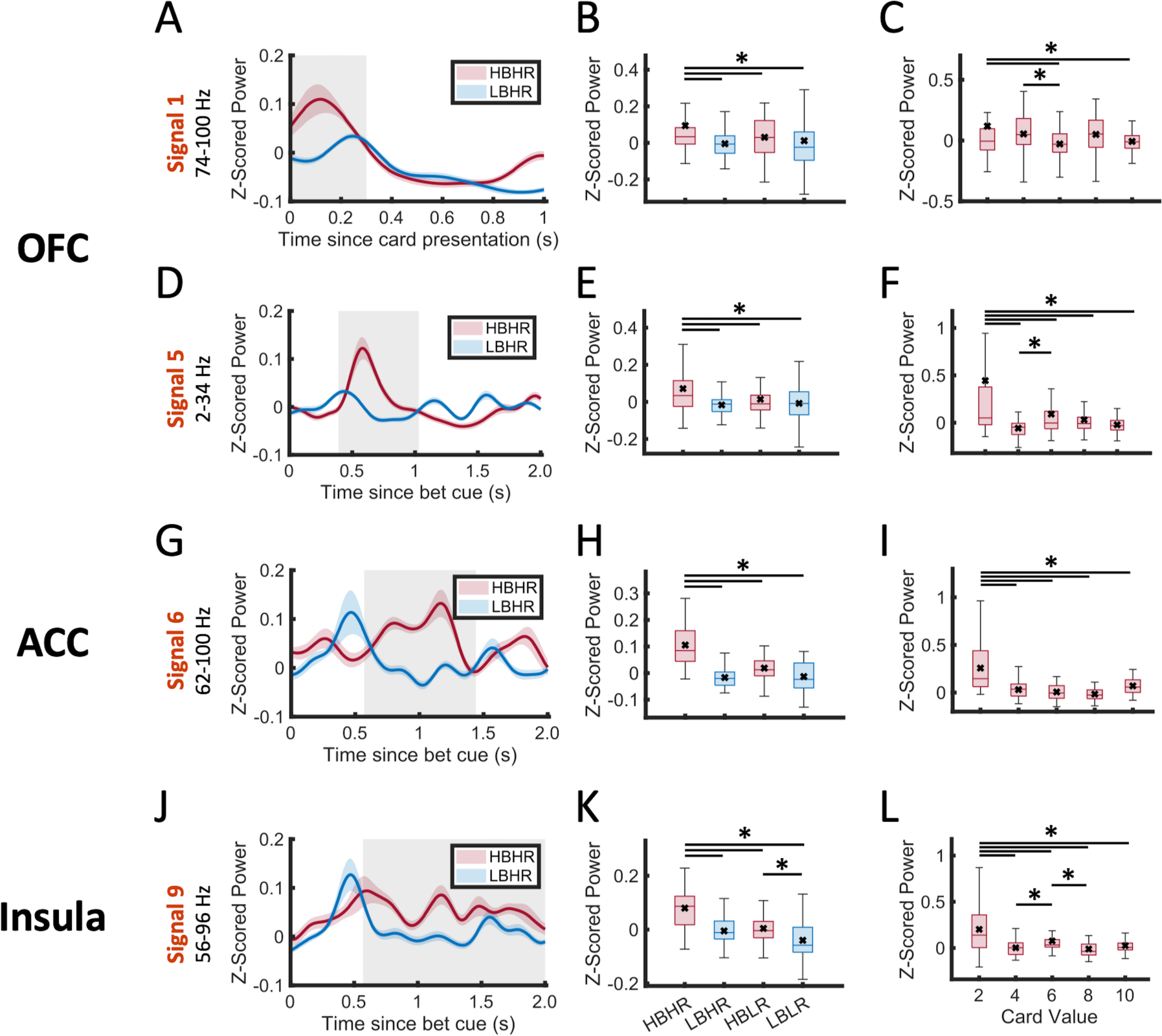
Risky decision signals. Average z-score power within cluster frequency range plotted over time for different conditions **(A,D,G,J)**. Shaded bars indicated standard error. Grey box indicates time range of significant clusters. Box plots of the mean power within the significant clusters across the four conditions **(B,E,H,K)**. Box plot of the mean power within the significant clusters prior to high bets for each patient card value **(C,F,I,L)**. Significant difference between groups (adjusted p<0.05) is indicated with line and * above. Black x indicates mean, whereas horizontal line within box indicates median. Orbitofrontal cortex signal 1 **(A-C)**, orbitofrontal cortex signal 5 **(D-F)**, anterior cingulate signal 6 **(G-I)**, Insula signal 9 (**J-L)**. HB = high bet; LB = low bet; HR = high risk; LR = low risk; OFC = orbitofrontal cortex, ACC = anterior cingulate cortex.

The ACC had one time-frequency cluster whose oscillatory power was associated with subsequent risky decisions. This signal had increased power in the gamma frequency range (62-100Hz) and occurred 0.58-1.45 seconds after bet cue presentation (*p*=4.0E-4 for cluster, *F*(1,142)=28.14 and adjusted *p*=4.5E-5 for interaction of bet and risk) (**Figures 3D, 5G,H**). The mean power within this cluster was highest in high risk trials in which patients subsequently bet high (adjusted *p*<0.0001 against all other risk-bet conditions) (**Figure 5G,H)**. Average power within this cluster was also significantly associated with both card value and the bet-card interaction (*F*(4,355)=23.5 and *F*(4,355)=16.6 respectively, both adjusted *p*<0.0001) (**Figure 5I**), with higher power prior to high bets for the patient card value of 2 (highest risk) compared to all other card values, again suggesting that this signal was modulated by risk level. Similarly, in insula there was a time frequency cluster associated with risky decisions which had increased power within the gamma frequency range (56-96 Hz) 0.58-2.01 seconds after bet cue presentation (*p*=0.0014 for cluster, *F*(1,306)=6.5 and adjusted *p*=0.013 for interaction of bet and risk) (**Figures 3H, 5J,K).** Power in this signal was significantly greater during high risk trials in which patients subsequently bet high compared to all other conditions (adjusted *p*<0.0001) (**Figure 5 J,K).** The power in this signal was also significantly associated with card value and the bet-card interaction (adjusted *p*<0.0001) (**Figure 5L**), and again was higher prior to high bets when the subject card was 2 compared to all other card values (adjusted *p*<0.0001).

### Optimized Decision Signals

The OFC, amygdala, and insula all had clusters in which z-scored power was significantly associated with subsequent decisions that optimized risk benefit ratio, in other words betting low in high-risk and high in low-risk conditions (cluster statistics *p*=2.0E-4, *p*=2.0E-4, *p*=4.0E-4 for each respectively) (**Figures 3A,E,G**). In the OFC, the time frequency cluster that was associated with a subsequent optimized decision was an increase in 2-26Hz power prior to optimal decisions that occurred 0.37-1.0 seconds after initial card presentation (**Figures 3A**, **6A,B**). The amygdala had a broad 2-96 Hz decrease in power prior to optimal decisions that occurred 0.07-1.00 seconds after initial card presentation (**Figures 3E**, **6C,D**). Similarly, the insula had a decrease in 2-60Hz power prior to optimized decisions that occurred 0.15-1.00 seconds after initial card presentation (**Figures 3G**, **6E,F**). Each of these signals had greater power in low risk scenarios when the patient subsequently bet high (all adjusted: *p*=0.022, *p*=3.7E-5, *p*=6.2E-5 for OFC, amygdala, and insula, respectively) and in high risk scenarios when the patient subsequently bet low (all adjusted: *p*=2.3E-19, *p*=2.6E-7, *p*=1.4E-11 respectively) on post-hoc testing (**Figure 6 B,D,F**).

**Figure 6.**
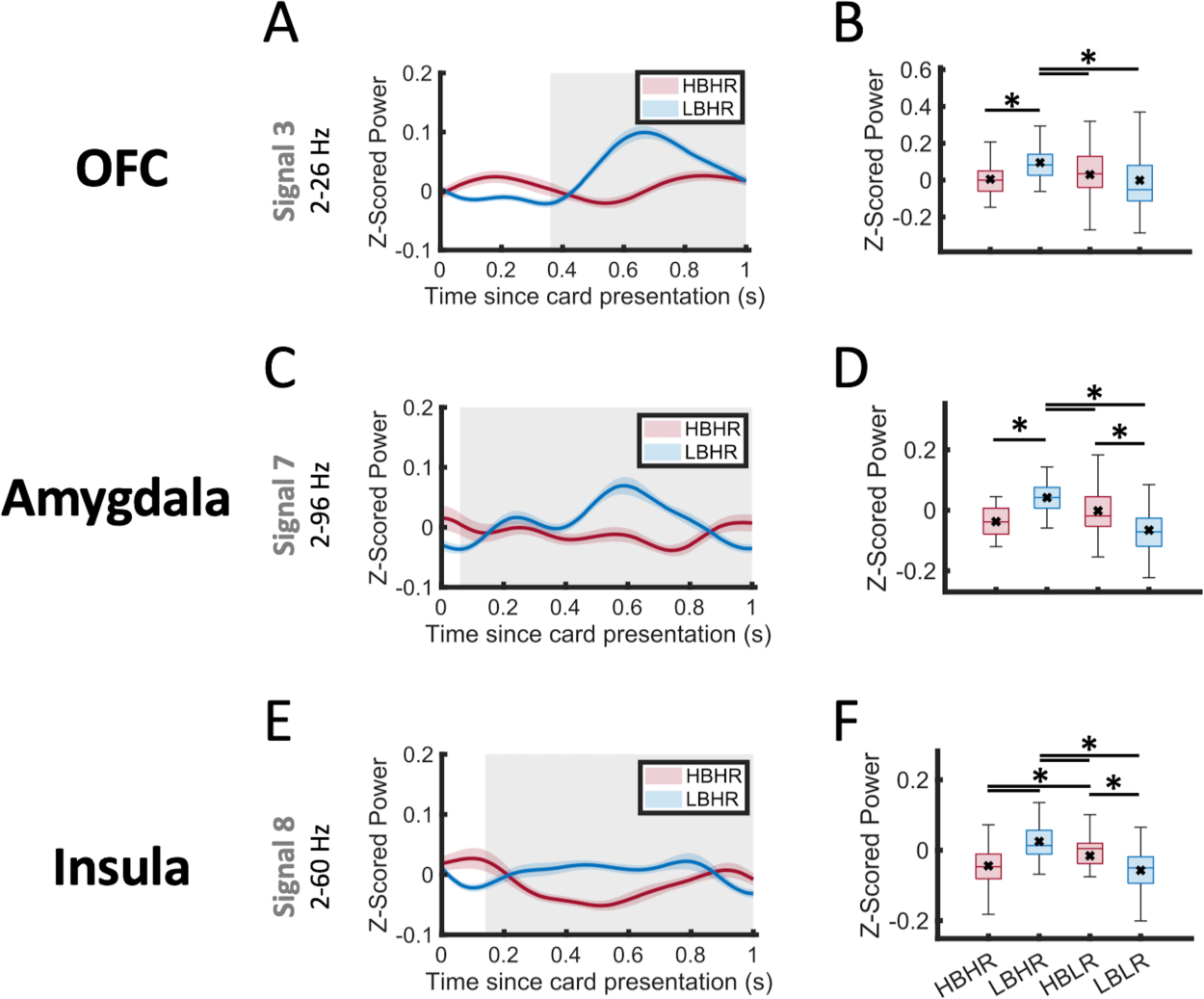
Optimized decision signals. **(A,C,E)** Average z-score power within cluster frequency range plotted over time for different conditions. Shaded bars indicate standard error. Grey boxes indicate time range of significant clusters. **(B,D,F)** Box plots of mean power within the significant clusters across the four conditions. Significant difference between groups (adjusted p<0.05) is indicated with line and * above. Black x indicates mean, whereas horizontal line within box indicates median. Orbitofrontal cortex (OFC) signal 3 **(A,B)**, amygdala signal 7 **(C,D)**, insula signal 8 **(E,F)**. (HB = high bet; LB = low bet; HR = high risk; LR = low risk).

### Relationship between Risk Signals and Impulsivity

We evaluated whether the signals that we had identified as predictive of risk-taking behavior were associated with a measure of overall impulsivity, which is associated with risk taking behavior. We computed difference in z-scored power within risky decision clusters between high and low bet conditions, and used linear regression models to evaluate the relationship between these values and BIS-11 scores. We found that insula risky decision signal power (**Figure 3H**) was significantly associated with BIS-11 scores (*p*=0.00478), with a smaller difference between power in high-bet and low-bet during high risk scenarios associated with greater impulsivity as measured by BIS-11 scores (adjusted *R*^2^=0.66) (**Figure 7A**). This indicates that patients with a more similar signal activity prior to betting high or low in a high-risk scenario were more impulsive. No significant relationship was found between the three other risky decision signals and BIS-11 scores.

**Figure 7.**
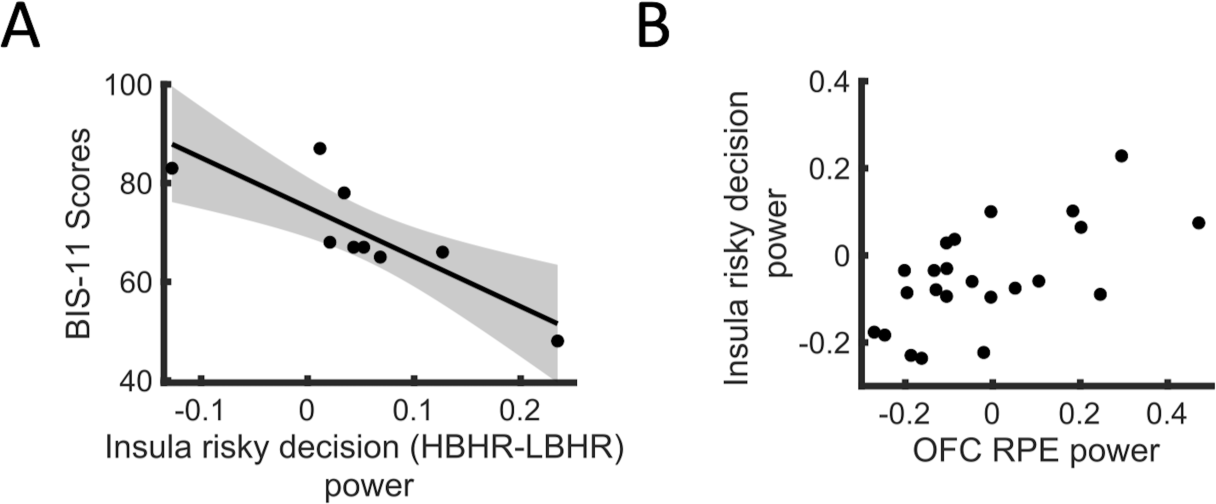
Relationships of insula risky decision signal to impulsivity and reward prediction error signals. **(A)** Scatterplot of the average difference in insula cluster (Signal 8 in Figure 3) z-scored power between high and low bet decisions in high risk trials compared against patient Barratt Impulsiveness Scale scores. Linear regression model plotted (estimated coefficient = −100.45, adjusted R^2^ = 0.66, p = 0.00478) with 95% confidence intervals shaded in grey. **(B)** Scatterplot of insula risky decision signal average Z-scored power (Signal 8 in Figure 3) association with the average Z-scored power of the high frequency OFC reward prediction error signal. (p=0.0048) HBHR = high bet high risk; LBHR = low risk high bet, RPE = reward prediction error, BIS-11 = Barratt Impulsiveness Scale.

### Relationship between Risk and Reward Signals

Finally, to evaluate the relationship between risk and reward signalling we used linear mixed effects models to determine whether the 4 identified RPE signals were associated with each identified risky decision signal. In a well-fitted model (adjusted *R*^2^=0.75), an increase in high frequency OFC RPE signal power was a significant predictor of increased insula risky decision signal power (*p*=0.0048) (**Figure 7B**, **Table 1**). The other three RPE signals in this model were not significantly associated (all three *p*>0.0125) (**Table 1**).

**Table 1.** Linear mixed effect models that predict risky decision signal power. R^2^, p values, and coefficients for each linear mixed effects model predicting risky decision signal mean power from reward prediction error (RPE) signals’ mean power. OFC = orbitofrontal cortex; RPE = reward prediction error; Coef=coefficient; RPE = reward prediction error signal

| Risky Decision Signals | Adjusted R<sup>2</sup> | RPE Signals |  |  |  |
| --- | --- | --- | --- | --- | --- |
|  |  | OFC (low frequency) | OFC (high frequency) | Anterior Cingulate | Amygdala |
| OFC Signal 1 | 0.082 | p=0.59<br>Coef= -.036 | p = .90<br>Coef= .014 | p = .60<br>Coef = -.05 | p = .45<br>Coef = .09 |
| OFC Signal 5 | 0.079 | p = .29<br>Coef = -.12 | p = .070<br>Coef = -.34 | p = .67<br>Coef = .067 | p = .66<br>Coef = .08 |
| Anterior Cingulate Signal 6 | 0.72 | p = .087<br>Coef = -.11 | p = .28<br>Coef = .12 | p = .48<br>Coef = .07 | p = .013<br>Coef = .31 |
| Insula Signal 9 | 0.75 | p = .21<br>Coef = -.06 | **p = .0048**<br>**Coef = .26** | p = 0.04<br>Coef = .16 | p = .021<br>Coef=.20 |

Bold indicates $P < 0.0125$ .

## Discussion

In this study we identify and characterize risk and reward related oscillatory power signals in the OFC, amygdala, ACC, and insula. We identify power changes associated with betting high in high-risk conditions in OFC, ACC, and insula that occur prior to a response being made, suggesting that these signals may be predictive of a subsequent risky decision. In the insula, the difference in this signal between subsequent bet choices in high-risk conditions was inversely associated with impulsivity scores. We also identified RPE signals in amygdala, ACC, and OFC, and found that a high frequency OFC RPE signal was correlated with insula risky decision signalling. Together these findings deepen our understanding of how reward circuitry signalling may influence risk taking.

### Reward Prediction Error Signals

We identified several signals associated with RPE. In ACC, amygdala, and OFC there were gamma range power increases with unexpected rewards, and in OFC there was also a decrease in low frequency power with unexpected rewards. This aligns with prior literature which has identified roles of the ACC, amygdala, and OFC in RPE signalling. Human electrophysiology studies have shown ventromedial prefrontal cortex and medial prefrontal cortex, including ACC, have alpha-band event related potential amplitudes associated with positive RPE.^47^ ACC has also been shown to have an important role in signalling negative outcomes.^48–50^ While we did not examine loss related activity in the present study, our findings confirm the role of ACC in signalling positive RPE. The OFC has also been found to play a role in RPE signalling,^51^ with some controversy existing on whether it plays an indirect role, signalling purely reward value,^52^ versus a direct role specifically signalling RPE-related information dependent on reward expectation.^53–57^ Prior studies supporting the indirect role of the OFC suggest that it does not specifically record an RPE signal, but records value and decision variables (such as probability) separately.^58^ Our findings support a direct role of the OFC in RPE signalling, as we find signals that differ in activity in unexpected reward from expected reward: one a lower frequency signal that is also modulated by reward magnitude, and the other a higher frequency signal whose power association with reward expectation is independent of reward magnitude. The amygdala is well-known to be strongly associated with reward outcome and reward evaluation^51,59^ and prior fMRI research has suggested that it has activity changes with both positive and negative RPEs.^60^ While several previous studies have identified positive RPE signals in the insula using both fMRI and intracranial EEG,^51,61^ we did not identify insula RPE signals in our study. This could be due to spatial sampling of the insula in our study, as different insula regions have been previously been proposed to play different roles in reward signalling.^62^

### Risk Encoding

We identified a gamma range signal following bet cue presentation in OFC that increased in power with increasing risk but was not associated with bet choice in the high-risk condition. This finding reinforces those of prior studies that have identified a role for OFC in risk signalling. Primate studies have shown that OFC single neuron activity encodes risk separate from value.^63^ Moreover, a previous study in human subjects found that OFC local field potentials are modulated by both reward probability evaluation, varying with chance of receiving a reward, and by uncertainty of reward, with maximal modulation in the most uncertain condition.^64^ Interestingly, in this prior study the reward probability signals that varied with percent chance of winning, which we in this study refer to as “risk” signals, occurred 400-600ms after cue presentation; the signals associated with uncertainty occurred 1000ms-2000ms after cue presentation. This timing is similar to the risk signal identified in our study, which occurred 230-1480ms after bet cue presentation.

### Risk Associated Decision Signals

We identified several power changes in the reward circuit structures examined that predicted optimal decision making under different risk conditions. In OFC, amygdala, and insula we identified specific power changes prior to decisions that maximized gains and minimized losses. These signals associated with subsequent optimized choices imply a role for these regions in decision-making and strategy. It is well-known that the prefrontal cortex is responsible for many aspects of decision-making.^65,66^ The OFC in particular is known to play a role in influencing decision-making based on prior outcomes,^67,68^ with decreased OFC activation associated with impairments in decision-making.^69^ Additionally, nonhuman primate studies have suggested that the amygdala plays a role both in encoding and in comparing values of presented objects prior to decision-making.^70^ Our study supports this role of the amygdala in optimal decision-making as we found an overall increase in broadband power prior to making a decision that maximized earnings according to the card’s associated risk level. Prior literature has also suggested that individuals with lesions of the anterior insula may have impairments in making decisions about how long to wait for an uncertain reward, suggesting that the insula is important in weighing possible rewards and costs.^71^ The anterior insula may also be important in social decision making, with fMRI activation in this structure associated with strategic decisions that maximize personal gain and minimize chance of rejection by a partner in social settings.^72^ Supporting the findings of these prior studies, we identify neurophysiological signals within the OFC, amygdala, and insula that are associated with decisions that maximize gain and minimize loss, suggesting a role for these regions in reward and punishment based decision making.

Neural activity associated with risk-taking has been previously demonstrated in reward regions including OFC, ACC, and insula, and our study reinforces and expands upon these findings. In OFC, we identified a high gamma frequency power increase during viewing of cues as well as a lower frequency power increase following bet cue that preceded risky decisions. Moreover, we also identified a broad gamma power decrease later on during cue presentation that preceded decisions to avoided risk. Previous studies have shown that resting state functional connectivity between OFC and inferior frontal gyrus is associated with risk-taking, and that individuals with internet gaming disorder have increased functional connectivity between these regions.^73^ Another fMRI study found that OFC activity and functional connectivity to ACC and ventromedial prefrontal cortices were associated with greater overall risk-taking tendency.^30^ Animal studies have suggested an important role for dopaminergic signalling in OFC in risk taking, with blockade of D1 OFC dopamine receptors decreasing risky decisions while blockade of D2 OFC dopamine receptors increases subsequent risky decisions.^74^ Building on these previous findings, we demonstrate specific OFC neurophysiology changes that precede decisions to take and avoid risk. Our findings suggest that OFC plays a role in signalling risky decisions through increased oscillatory activity across frequencies, but can also signal risk aversion through decreased oscillatory activity. Prior literature in nonhuman primates suggests that OFC neural firing represents the value of choices during decision making and may provide this value information to ACC to guide decision-making and selection of response.^75^ Interestingly, we also identified an ACC gamma power increase predictive of a subsequent decision to take a risk that occurred immediately after the OFC risk-related signal. Though we did not investigate associations between these two signals, this similarity in timing and frequency may be of interest for future investigation.

Lastly, we identified an insula gamma power increase following bet cue that predicted subsequent risky choices. The insula is broadly thought to be implicated in risk related decision making and reward-seeking,^76,77^ and decreased insula fMRI activity has been associated with higher levels of risk aversion in patients with obsessive-compulsive disorder.^26^ Our study supports the role of insula in risk related decision-making and suggests that insula neural signals can predict risky choices.

### Risky Decision Signal Associations with Impulsivity and Reward Prediction Error

Interestingly, we found a relationship between insula risky decision signal and impulsivity. We found that a smaller difference in insula gamma power prior to high versus low bets in high risk conditions was associated with greater impulsivity as measured by BIS-11 scores. Previously, neuroimaging studies have suggested that increased insula thickness is associated with increased impulsivity^78^ and that increased insula activity during reward and loss anticipation is associated with greater impulsivity as measured by personality inventories.^79^ Our findings expand on this by suggesting that individuals with higher impulsivity have less differentiation in neural representation of high and low risk choices. To our knowledge, the electrophysiology of this insula relationship with impulsivity has not been previously elucidated.

Moreover, we also find that this insula risky decision signal is positively correlated with OFC RPE power. Though the insula and OFC specifically have not previously been implicated in this interaction between RPE and risk-taking decisions, this relationship has been suggested in other brain regions using other methodological techniques. For example, prior event-related potential studies using scalp EEG have demonstrated that frontal feedback related negativity signal amplitudes may be associated with the level of risk taken, with reduced amplitudes in high-risk choices.^80^ Additionally, neuroeconomic studies in primates have suggested that dopaminergic RPE signals are associated with an economic representation of subjective value at the time of decision.^81^ Imaging studies suggest that risk prediction error signalling in the inferior frontal gyrus may be more pronounced in risk averse patients.^82^ In the nucleus accumbens, a relationship between dopaminergic cells signalling unfavourable outcomes has been shown to play a role in real-time risky decision making.^83^ However, to our knowledge, the relationship as described here between OFC post-outcome RPE signal and pre-decision insula risky decision signal has not before been described. Future research may be necessary to better examine the elements of these relationships between risky decision signals and RPE signals, as they may be modulated by learning, magnitude of risk and reward, or other factors.

### Limitations

This is a single-institution study with a specialized population of individuals with medically resistant epilepsy. Epilepsy may alter neural activity patterns, though we excluded electrode contacts within the seizure onset zone from analysis to decrease risk of this confounder. Furthermore, despite a homogenous patient population, the average BIS-11 score was 66.9 +/− 10.0, which is comparable to a previously reported adult average of 62.3 +/− 10.3,^38^ suggesting similar impulsivity to the general population. Additionally, cluster-based permutation is not suitable to identify effect latencies on a millisecond scale; therefore timing and frequency of reported signals are approximate.^84^ Notably, this experimental paradigm also did not have any option to entirely avoid risk; patients were required to place a bet. Future studies may implement levels of risk with an option to avoid risk entirely to assess these regions given more complex options.

## Data availability

The data that support the findings of this study are available from the corresponding author upon reasonable request.

## Supplementary material

Supplementary material is available at *Brain* online.

## References

1. Lane SD, Cherek DR. Analysis of risk taking in adults with a history of high risk behavior. Drug and Alcohol Dependence. 2000;60(2):179–187. doi:10.1016/S0376-8716(99)00155-6

2. Bechara A, Damasio H. Decision-making and addiction (part I): impaired activation of somatic states in substance dependent individuals when pondering decisions with negative future consequences. Neuropsychologia. 2002;40(10):1675–1689. doi:10.1016/s0028-3932(02)00015-5

3. Frank D, Elliott L, Cleland CM, et al. “As safe as possible”: a qualitative study of opioid withdrawal and risk behavior among people who use illegal opioids. Harm Reduct J. 2023;20(1):158. doi:10.1186/s12954-023-00893-9

4. Chan CC, Alter S, Hazlett EA, et al. Neural correlates of impulsivity in bipolar disorder: A systematic review and clinical implications. Neurosci Biobehav Rev. 2023;147:105109. doi:10.1016/j.neubiorev.2023.105109

5. Boutoleau-Bretonnière C, Kapogiannis D, El Haj M. The guaranteed euros: Probabilistic discounting in behavioural-variant frontotemporal dementia. J Neuropsychol. Published online December 22, 2023. doi:10.1111/jnp.12357

6. Colautti L, Iannello P, Silveri MC, Antonietti A. Decision making in Parkinson’s disease: An analysis of the studies using the Iowa Gambling Task. Eur J Neurosci. 2021;54(10):7513–7549. doi:10.1111/ejn.15497

7. Evens R, Hoefler M, Biber K, Lueken U. The Iowa Gambling Task in Parkinson’s disease: A meta-analysis on effects of disease and medication. Neuropsychologia. 2016;91:163–172. doi:10.1016/j.neuropsychologia.2016.07.032

8. Kjær SW, Damholdt MF, Callesen MB. A systematic review of decision-making impairments in Parkinson’s Disease: Dopaminergic medication and methodological variability. Basal Ganglia. 2018;14:31–40. doi:10.1016/j.baga.2018.07.003

9. Smulders K, Esselink RA, Cools R, Bloem BR. Trait Impulsivity Is Associated with the Risk of Falls in Parkinson’s Disease. PLoS One. 2014;9(3):e91190. doi:10.1371/journal.pone.0091190

10. Karlsson A, Håkansson A. Gambling disorder, increased mortality, suicidality, and associated comorbidity: A longitudinal nationwide register study. J Behav Addict. 7(4):1091–1099. doi:10.1556/2006.7.2018.112

11. Blonigen DM, Timko C, Moos BS, Moos RH. Impulsivity is an Independent Predictor of 15-Year Mortality Risk among Individuals Seeking Help for Alcohol-Related Problems. Alcohol Clin Exp Res. 2011;35(11):2082–2092. doi:10.1111/j.1530-0277.2011.01560.x

12. Voon V, Fox SH. Medication-related impulse control and repetitive behaviors in Parkinson disease. Arch Neurol. 2007;64(8):1089–1096. doi:10.1001/archneur.64.8.1089

13. Loya JM, Benitez B, Kiluk BD. The Effect of Cognitive Behavioral Therapy on Impulsivity in Addictive Disorders: a Narrative Review. Curr Addict Rep. 2023;10(3):485–493. doi:10.1007/s40429-023-00491-6

14. Sethi K. Levodopa unresponsive symptoms in Parkinson disease. Movement Disorders. 2008;23(S3):S521–S533. doi:10.1002/mds.22049

15. Probst CC, van Eimeren T. The Functional Anatomy of Impulse Control Disorders. Curr Neurol Neurosci Rep. 2013;13(10):386. doi:10.1007/s11910-013-0386-8

16. Rolls ET, Cheng W, Feng J. The orbitofrontal cortex: reward, emotion and depression. Brain Commun. 2020;2(2):fcaa196. doi:10.1093/braincomms/fcaa196

17. Dolan R j. The human amygdala and orbital prefrontal cortex in behavioural regulation. Philosophical Transactions of the Royal Society B: Biological Sciences. 2007;362(1481):787–799. doi:10.1098/rstb.2007.2088

18. Cai J, Tong Q. Anatomy and Function of Ventral Tegmental Area Glutamate Neurons. Front Neural Circuits. 2022;16:867053. doi:10.3389/fncir.2022.867053

19. Lewis RG, Florio E, Punzo D, Borrelli E. The Brain’s Reward System in Health and Disease. Adv Exp Med Biol. 2021;1344:57–69. doi:10.1007/978-3-030-81147-1_4

20. Morales M, Margolis EB. Ventral tegmental area: cellular heterogeneity, connectivity and behaviour. Nat Rev Neurosci. 2017;18(2):73–85. doi:10.1038/nrn.2016.165

21. Freeman C, Dirks M, Weinberg A. Neural response to rewards predicts risk-taking in late but not early adolescent females. Dev Cogn Neurosci. 2020;45:100808. doi:10.1016/j.dcn.2020.100808

22. Rutledge RB, Smittenaar P, Zeidman P, et al. Risk Taking for Potential Reward Decreases across the Lifespan. Current Biology. 2016;26(12):1634–1639. doi:10.1016/j.cub.2016.05.017

23. Moeller M, Grohn J, Manohar S, Bogacz R. An association between prediction errors and risk-seeking: Theory and behavioral evidence. PLoS Comput Biol. 2021;17(7):e1009213. doi:10.1371/journal.pcbi.1009213

24. Stopper CM, Tse MTL, Montes DR, Wiedman CR, Floresco SB. Overriding Phasic Dopamine Signals Redirects Action Selection during Risk/Reward Decision Making. Neuron. 2014;84(1):177–189. doi:10.1016/j.neuron.2014.08.033

25. Tahmasian M, Rochhausen L, Maier F, et al. Impulsivity is Associated with Increased Metabolism in the Fronto-Insular Network in Parkinson’s Disease. Frontiers in Behavioral Neuroscience. 2015;9. Accessed February 26, 2024. https://www.frontiersin.org/articles/10.3389/fnbeh.2015.00317

26. Han Y, Gao F, Wang X, et al. Neural correlates of risk taking in patients with obsessive-compulsive disorder during risky decision-making. J Affect Disord. 2024;345:192–199. doi:10.1016/j.jad.2023.10.099

27. Mao T, Fang Z, Chai Y, et al. Sleep deprivation attenuates neural responses to outcomes from risky decision-making. Psychophysiology. Published online October 31, 2023:e14465. doi:10.1111/psyp.14465

28. Wang L, Zheng H, Wang M, Chen S, Du X, Dong GH. Sex differences in neural substrates of risk taking: Implications for sex-specific vulnerabilities to internet gaming disorder. J Behav Addict. 2022;11(3):778–795. doi:10.1556/2006.2022.00057

29. Woo JH, Azab H, Jahn A, Hayden B, Brown JW. The PRO model accounts for the anterior cingulate cortex role in risky decision-making and monitoring. Cogn Affect Behav Neurosci. 2022;22(5):952–968. doi:10.3758/s13415-022-00992-3

30. Rolls ET, Wan Z, Cheng W, Feng J. Risk-taking in humans and the medial orbitofrontal cortex reward system. Neuroimage. 2022;249:118893. doi:10.1016/j.neuroimage.2022.118893

31. Ren P, Ma M, Zhuang Y, et al. Dorsal and ventral fronto-amygdala networks underlie risky decision-making in age-related cognitive decline. Geroscience. Published online September 12, 2023. doi:10.1007/s11357-023-00922-2

32. Brevers D, Baeken C, Bechara A, et al. Increased brain reactivity to gambling unavailability as a marker of problem gambling. Addict Biol. 2021;26(4):e12996. doi:10.1111/adb.12996

33. Poudel R, Tobia MJ, Riedel MC, et al. Risky decision-making strategies mediate the relationship between amygdala activity and real-world financial savings among individuals from lower income households: A pilot study. Behav Brain Res. 2022;428:113867. doi:10.1016/j.bbr.2022.113867

34. Glover GH. Overview of Functional Magnetic Resonance Imaging. Neurosurg Clin N Am. 2011;22(2):133–139. doi:10.1016/j.nec.2010.11.001

35. Studer B, Pedroni A, Rieskamp J. Predicting Risk-Taking Behavior from Prefrontal Resting-State Activity and Personality. PLoS One. 2013;8(10):e76861. doi:10.1371/journal.pone.0076861

36. Dantas AM, Sack AT, Bruggen E, Jiao P, Schuhmann T. Reduced risk-taking behavior during frontal oscillatory theta band neurostimulation. Brain Research. 2021;1759:147365. doi:10.1016/j.brainres.2021.147365

37. Polezzi D, Sartori G, Rumiati R, Vidotto G, Daum I. Brain correlates of risky decision-making. NeuroImage. 2010;49(2):1886–1894. doi:10.1016/j.neuroimage.2009.08.068

38. Stanford MS, Mathias CW, Dougherty DM, Lake SL, Anderson NE, Patton JH. Fifty years of the Barratt Impulsiveness Scale: An update and review. Personality and Individual Differences. 2009;47(5):385–395. doi:10.1016/j.paid.2009.04.008

39. D’Haese PF, Pallavaram S, Li R, et al. CranialVault and its CRAVE tools: a clinical computer assistance system for deep brain stimulation (DBS) therapy. Med Image Anal. 2012;16(3):744–753. doi:10.1016/j.media.2010.07.009

40. Desikan RS, Ségonne F, Fischl B, et al. An automated labeling system for subdividing the human cerebral cortex on MRI scans into gyral based regions of interest. Neuroimage. 2006;31(3):968–980. doi:10.1016/j.neuroimage.2006.01.021

41. Fischl B, Salat DH, Busa E, et al. Whole brain segmentation: automated labeling of neuroanatomical structures in the human brain. Neuron. 2002;33(3):341–355. doi:10.1016/s0896-6273(02)00569-x

42. Fischl B, Salat DH, van der Kouwe AJW, et al. Sequence-independent segmentation of magnetic resonance images. Neuroimage. 2004;23 Suppl 1:S69–84. doi:10.1016/j.neuroimage.2004.07.016

43. Brainard DH. The Psychophysics Toolbox. Spat Vis. 1997;10(4):433–436.

44. Singh B, Wang Z, Madiah LM, et al. Brain-wide human oscillatory local field potential activity during visual working memory. iScience. 2024;27(3):109130. doi:10.1016/j.isci.2024.109130

45. Oostenveld R, Fries P, Maris E, Schoffelen JM. FieldTrip: Open source software for advanced analysis of MEG, EEG, and invasive electrophysiological data. Comput Intell Neurosci. 2011;2011:156869. doi:10.1155/2011/156869

46. Maris E, Oostenveld R. Nonparametric statistical testing of EEG- and MEG-data. J Neurosci Methods. 2007;164(1):177–190. doi:10.1016/j.jneumeth.2007.03.024

47. Oya H, Adolphs R, Kawasaki H, Bechara A, Damasio A, Howard MA. Electrophysiological correlates of reward prediction error recorded in the human prefrontal cortex. Proceedings of the National Academy of Sciences. 2005;102(23):8351–8356. doi:10.1073/pnas.0500899102

48. Gehring WJ, Willoughby AR. The medial frontal cortex and the rapid processing of monetary gains and losses. Science. 2002;295(5563):2279–2282. doi:10.1126/science.1066893

49. Miltner WHR, Lemke U, Weiss T, Holroyd C, Scheffers MK, Coles MGH. Implementation of error-processing in the human anterior cingulate cortex: a source analysis of the magnetic equivalent of the error-related negativity. Biol Psychol. 2003;64(1-2):157–166. doi:10.1016/s0301-0511(03)00107-8

50. Yeung N, Holroyd CB, Cohen JD. ERP correlates of feedback and reward processing in the presence and absence of response choice. Cereb Cortex. 2005;15(5):535–544. doi:10.1093/cercor/bhh153

51. Ulrich M, Rüger A, Durner V, Grön G, Graf H. Reward is not reward: Differential impacts of primary and secondary rewards on expectation, outcome, and prediction error in the human brain’s reward processing regions. NeuroImage. 2023;283:120440. doi:10.1016/j.neuroimage.2023.120440

52. Stalnaker TA, Liu TL, Takahashi YK, Schoenbaum G. Orbitofrontal neurons signal reward predictions, not reward prediction errors. Neurobiology of Learning and Memory. 2018;153:137–143. doi:10.1016/j.nlm.2018.01.013

53. Knutson B, Wimmer GE. Splitting the Difference. Annals of the New York Academy of Sciences. 2007;1104(1):54–69. doi:10.1196/annals.1390.020

54. Nobre AC, Coull JT, Frith CD, Mesulam MM. Orbitofrontal cortex is activated during breaches of expectation in tasks of visual attention. Nat Neurosci. 1999;2(1):11–12. doi:10.1038/4513

55. O’Doherty JP, Dayan P, Friston K, Critchley H, Dolan RJ. Temporal Difference Models and Reward-Related Learning in the Human Brain. Neuron. 2003;38(2):329–337. doi:10.1016/S0896-6273(03)00169-7

56. Sul JH, Kim H, Huh N, Lee D, Jung MW. Distinct Roles of Rodent Orbitofrontal and Medial Prefrontal Cortex in Decision Making. Neuron. 2010;66(3):449–460. doi:10.1016/j.neuron.2010.03.033

57. Tobler PN, O’Doherty JP, Dolan RJ, Schultz W. Human Neural Learning Depends on Reward Prediction Errors in the Blocking Paradigm. Journal of Neurophysiology. 2006;95(1):301–310. doi:10.1152/jn.00762.2005

58. Wallis JD, Kennerley SW. Contrasting reward signals in the orbitofrontal cortex and anterior cingulate cortex. Annals of the New York Academy of Sciences. 2011;1239(1):33–42. doi:10.1111/j.1749-6632.2011.06277.x

59. Plastic and stimulus-specific coding of salient events in the central amygdala - PubMed. Accessed January 26, 2024. https://pubmed.ncbi.nlm.nih.gov/37020025/

60. Kolada E, Bielski K, Wilk M, et al. The Human Centromedial Amygdala Contributes to Negative Prediction Error Signaling during Appetitive and Aversive Pavlovian Gustatory Learning. J Neurosci. 2023;43(17):3176–3185. doi:10.1523/JNEUROSCI.0926-22.2023

61. Hoy CW, Quiroga-Martinez DR, Sandoval E, et al. Asymmetric coding of reward prediction errors in human insula and dorsomedial prefrontal cortex. Nat Commun. 2023;14(1):8520. doi:10.1038/s41467-023-44248-1

62. Castro DC, Berridge KC. Opioid and orexin hedonic hotspots in rat orbitofrontal cortex and insula. Proc Natl Acad Sci U S A. 2017;114(43):E9125–E9134. doi:10.1073/pnas.1705753114

63. Schultz W, O’Neill M, Tobler PN, Kobayashi S. Neuronal signals for reward risk in frontal cortex. Annals of the New York Academy of Sciences. 2011;1239(1):109–117. doi:10.1111/j.1749-6632.2011.06256.x

64. Li Y, Vanni-Mercier G, Isnard J, Mauguière F, Dreher JC. The neural dynamics of reward value and risk coding in the human orbitofrontal cortex. Brain. 2016;139(4):1295–1309. doi:10.1093/brain/awv409

65. Tobler PN, Christopoulos GI, O’Doherty JP, Dolan RJ, Schultz W. Risk-dependent reward value signal in human prefrontal cortex. Proceedings of the National Academy of Sciences. 2009;106(17):7185–7190. doi:10.1073/pnas.0809599106

66. Pushkarskaya H, Smithson M, Joseph JE, Corbly C, Levy I. Neural Correlates of Decision-Making Under Ambiguity and Conflict. Frontiers in Behavioral Neuroscience. 2015;9. Accessed January 28, 2024. https://www.frontiersin.org/articles/10.3389/fnbeh.2015.00325

67. Tegelbeckers J, Porter DB, Voss JL, Schoenbaum G, Kahnt T. Lateral orbitofrontal cortex integrates predictive information across multiple cues to guide behavior. Curr Biol. 2023;33(20):4496–4504.e5. doi:10.1016/j.cub.2023.09.033

68. Morgane PJ, Galler JR, Mokler DJ. A review of systems and networks of the limbic forebrain/limbic midbrain. Progress in Neurobiology. 2005;75(2):143–160. doi:10.1016/j.pneurobio.2005.01.001

69. Li Z, Zhang W, Du Y. Neural mechanisms of intertemporal and risky decision-making in individuals with internet use disorder: A perspective from directed functional connectivity. J Behav Addict. 2023;12(4):907–919. doi:10.1556/2006.2023.00068

70. Grabenhorst F, Ponce-Alvarez A, Battaglia-Mayer A, Deco G, Schultz W. A view-based decision mechanism for rewards in the primate amygdala. Neuron. 2023;111(23):3871–3884.e14. doi:10.1016/j.neuron.2023.08.024

71. van Geen C, Chen Y, Kazinka R, Vaidya AR, Kable JW, McGuire JT. Lesions to different regions of frontal cortex have dissociable effects on voluntary persistence. bioRxiv. Published online November 17, 2023:2023.11.16.567406. doi:10.1101/2023.11.16.567406

72. Sazhin D, Wyngaarden JB, Dennison JB, et al. Trait Reward Sensitivity Modulates Connectivity with the Temporoparietal Junction and Anterior Insula during Strategic Decision Making. bioRxiv. Published online October 19, 2023:2023.10.19.563125. doi:10.1101/2023.10.19.563125

73. Liu S, Lu Y, Li S, et al. Resting-state functional connectivity within orbitofrontal cortex and inferior frontal gyrus modulates the relationship between reflection level and risk-taking behavior in internet gaming disorder. Brain Res Bull. 2022;178:49–56. doi:10.1016/j.brainresbull.2021.10.019

74. Jenni NL, Li YT, Floresco SB. Medial orbitofrontal cortex dopamine D1/D2 receptors differentially modulate distinct forms of probabilistic decision-making. Neuropsychopharmacol. 2021;46(7):1240–1251. doi:10.1038/s41386-020-00931-1

75. Balewski ZZ, Elston TW, Knudsen EB, Wallis JD. Value dynamics affect choice preparation during decision-making. Nat Neurosci. 2023;26(9):1575–1583. doi:10.1038/s41593-023-01407-3

76. McGregor MS, LaLumiere RT. Still a “hidden island”? The rodent insular cortex in drug seeking, reward, and risk. Neurosci Biobehav Rev. 2023;153:105334. doi:10.1016/j.neubiorev.2023.105334

77. Zaleskiewicz T, Traczyk J, Sobkow A, Fulawka K, Megías-Robles A. Visualizing risky situations induces a stronger neural response in brain areas associated with mental imagery and emotions than visualizing non-risky situations. Front Hum Neurosci. 2023;17:1207364. doi:10.3389/fnhum.2023.1207364

78. Churchwell JC, Yurgelun-Todd DA. Age-related changes in insula cortical thickness and impulsivity: Significance for emotional development and decision-making. Dev Cogn Neurosci. 2013;6:80–86. doi:10.1016/j.dcn.2013.07.001

79. Villafuerte S, Heitzeg MM, Foley S, et al. Impulsiveness and Insula activation during reward anticipation are associated with genetic variants in GABRA2 in a family sample enriched for alcoholism. Mol Psychiatry. 2012;17(5):511–519. doi:10.1038/mp.2011.33

80. Schuermann B, Endrass T, Kathmann N. Neural correlates of feedback processing in decision-making under risk. Front Hum Neurosci. 2012;6:204. doi:10.3389/fnhum.2012.00204

81. Stauffer WR, Lak A, Schultz W. Dopamine reward prediction error responses reflect marginal utility. Curr Biol. 2014;24(21):2491–2500. doi:10.1016/j.cub.2014.08.064

82. d’Acremont M, Lu ZL, Li X, Van der Linden M, Bechara A. Neural correlates of risk prediction error during reinforcement learning in humans. Neuroimage. 2009;47(4):1929–1939. doi:10.1016/j.neuroimage.2009.04.096

83. Zalocusky KA, Ramakrishnan C, Lerner TN, Davidson TJ, Knutson B, Deisseroth K. Nucleus accumbens D2R cells signal prior outcomes and control risky decision-making. Nature. 2016;531(7596):642–646. doi:10.1038/nature17400

84. Sassenhagen J, Draschkow D. Cluster-based permutation tests of MEG/EEG data do not establish significance of effect latency or location. Psychophysiology. 2019;56(6):e13335. doi:10.1111/psyp.13335

