## Supplementary figures and images for "Reward Circuit Local Field Potential Modulations Precede Risk Taking"

### Supplemental Table 1

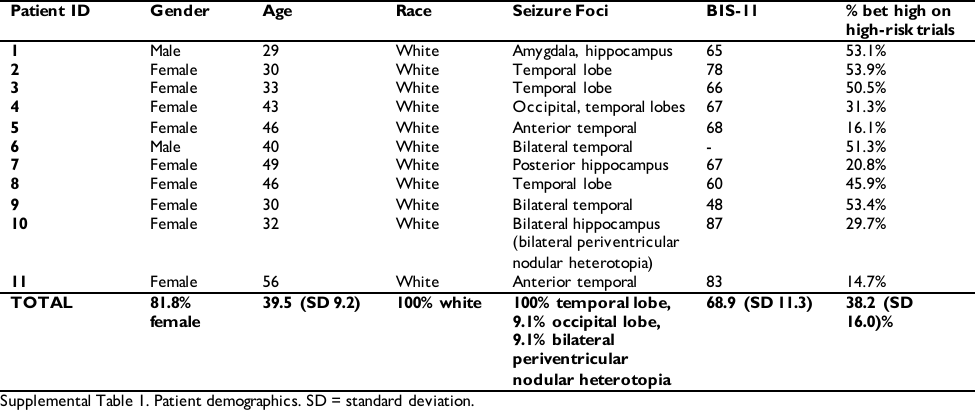
